# *Bordetella* Filamentous hemagglutinin (FhaB) and Adenylyl cyclase toxin (ACT) interactions on the bacterial surface are consistent with FhaB-mediated delivery of ACT to phagocytic cells

**DOI:** 10.1101/2023.09.06.556475

**Authors:** Zachary M. Nash, Carol S. Inatsuka, Peggy A. Cotter, Richard M. Johnson

## Abstract

*Bordetella* species that cause respiratory infections in mammals include *B. pertussis*, which causes human whooping cough, and *B. bronchiseptica*, which infects nearly all mammals. Both bacterial species produce filamentous hemagglutinin (FhaB) and adenylyl cyclase toxin (ACT), prominent surface-associated and secreted virulence factors that contribute to persistence in the lower respiratory tract by inhibiting clearance by phagocytic cells. FhaB and ACT proteins interact with themselves, each other, and host cells. Using immunoblot analyses, we showed that ACT binds to FhaB preferentially on the bacterial surface before being released into the extracellular environment. We showed that SphB1, a surface protease identified based on its ability to cleave FhaB, also cleaves ACT, and we showed that the presence of ACT blocks SphB1-dependent and independent cleavage of FhaB, but the presence of FhaB does not block SphB1-dependent cleavage of ACT. SphB1-dependent cleavage of ACT occurs proximally to ACT’s active site, and hence is predicted to inactivate ACT. We also showed that FhaB-bound ACT can intoxicate host cells producing CR3, the receptor for ACT. In addition to increasing our understanding of FhaB, ACT, and ACT interactions on the *Bordetella* surface, our data support a model in which FhaB functions as a novel toxin delivery system by binding to ACT and allowing its release upon binding of ACT to its receptor, CR3, on phagocytic cells.

**AUTHOR SUMMARY:** Bacteria need to control the variety, abundance, and conformation of proteins on the cellular surface to survive. Members of the Gram-negative bacterial genus *Bordetella* include *B. pertussis*, which causes whooping cough in humans, and *B. bronchiseptica*, which causes respiratory infections in a broad range of mammals. These extremely closely related species produce two prominent virulence factors, the two-partner secretion (TPS) effector FhaB and Adenylyl cyclase toxin (ACT), that interact with themselves, each other, and host cells. In this work, we showed that ACT binds preferentially to FhaB on the bacterial surface before being released into the extracellular environment. We showed that the exoprotease SphB1, which cleaves FhaB, also cleaves ACT. We showed that ACT that is bound to FhaB can be delivered to CR3^+^ host cells. Our data support a model in which FhaB functions to deliver ACT specifically to phagocytic cells, and not epithelial cells. This is the first report of a TPS system facilitating delivery of a separate polypeptide toxin to target cells and expands our understanding of how these systems contribute to bacterial pathogenesis.

## INTRODUCTION

Interactions between proteins on a bacterial surface with each other or with biotic or abiotic surfaces often have consequences that are critical to bacterial survival. Such interactions can mediate adherence of the bacteria to cells or surfaces and facilitate delivery of proteins, including toxins, to nearby cells. *Bordetella pertussis*, the causal agent of human whooping cough, and the closely related species *Bordetella bronchiseptica*, which causes respiratory infections in a broad range of mammals, produce two prominent virulence factors, filamentous hemagglutinin (FhaB) and adenylate cyclase toxin (ACT), that interact with themselves, each other(1), and molecules on host cells. Although both FhaB and ACT play roles in adherence(2–6), biofilm formation(7,8), and persistence in the lower respiratory tract(9–12), the mechanisms and importance of FhaB-ACT interactions in mediating these phenotypes are not well understood.

FhaB and its outer membrane transporter FhaC compose the prototypical two-partner secretion (TPS, also known as Type Vb) family of bacterial secretion systems that are broadly distributed among Gram-negative bacteria(13–15). FhaB is synthesized as a ∼375 kD preproprotein with an extended N-terminal signal sequence that mediates Sec-dependent delivery to the periplasm(16,17) and a TPS domain that interacts with FhaC’s periplasmic POTRA domains to initiate translocation across the outer membrane(18–20). According to the current model (Fig. 1), FhaB emerges on the bacterial surface as a hairpin with its N terminus anchored to FhaC while approximately two-thirds of FhaB are translocated through the FhaC barrel in the N- to C-terminal direction, forming a β-helical shaft topped with a globular mature C-terminal domain (MCD)(21). A ∼200 amino acid (aa) region, called the prodomain N terminus (PNT), forms a molecular knot that blocks translocation through FhaC such that the remaining C-terminal ∼1,200 aa of FhaB, called the prodomain (PD), are retained in the periplasm(22). Full-length FhaB forms a stable complex with FhaC until regulated processing of the PD begins with an unknown signal that promotes DegP-dependent removal of the extreme C terminus (ECT) of FhaB to form FhaB^CP^(23). The periplasmic protease CtpA is required for processive degradation of the remainder of the FhaB^CP^ PD to form a polypeptide called FHA’ (∼263 kD) that lacks a functional PNT and ultimately exits the FhaC channel(24). Under certain conditions, such as prolonged growth *in vitro*, the surface-anchored autotransporter protease SphB1 cleaves near the FHA’ C terminus, causing the immediate release of a ∼250 kD polypeptide called FHA. SphB1 can also cleave FhaB precursors at two alternative sites just N-terminal to the FHA cleavage site to form FHA_1_ and FHA_2_, and this activity is enhanced in the absence of CtpA(24). Although SphB1-dependent cleavage of FhaB has been reported to be a critical maturation process(25), the biological significance of SphB1 cleavage of FhaB is unclear.

**Figure 1.**
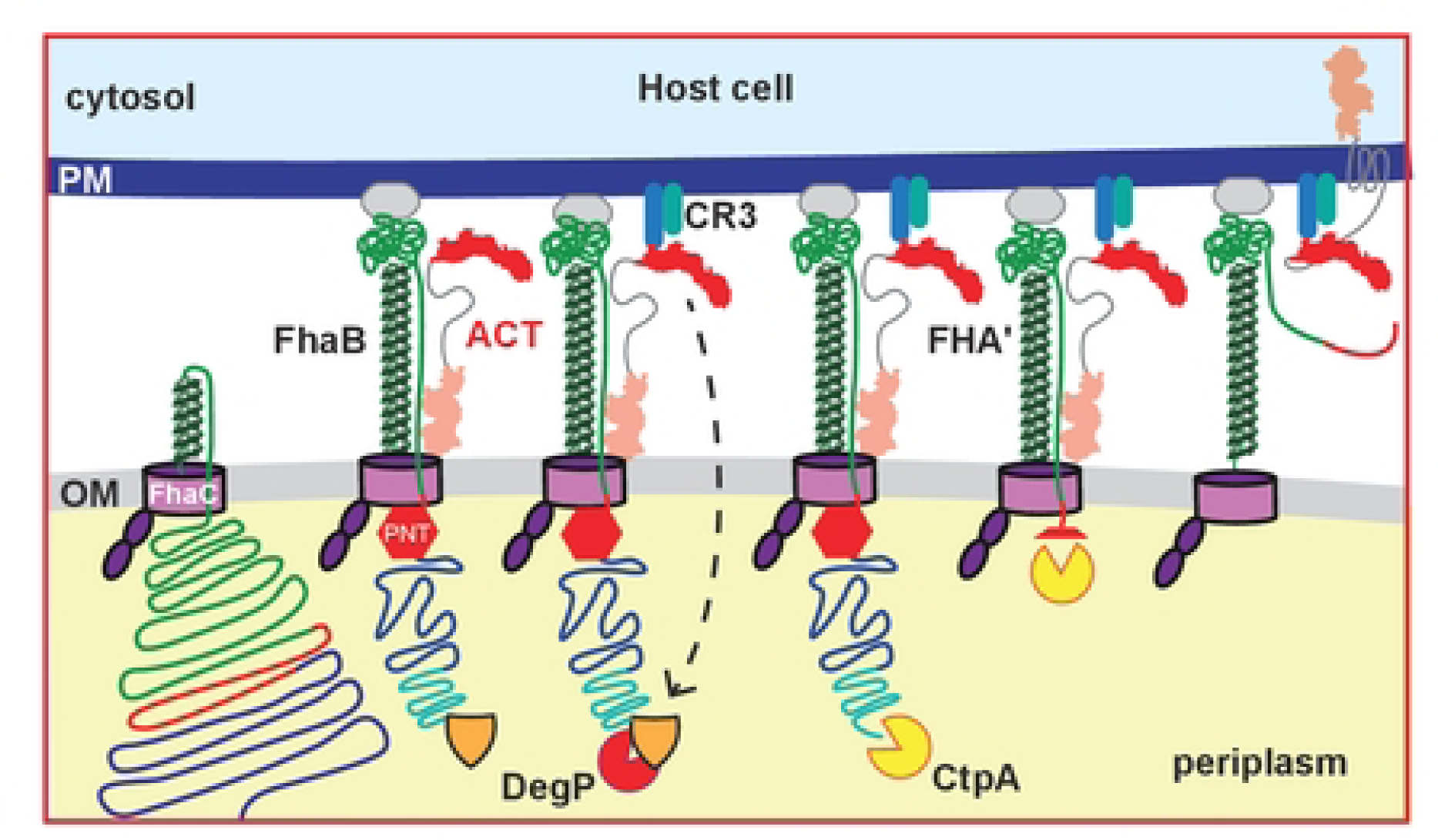
Hypothetical model for FhaB-mediated delivery of ACT to CR3+ cells. FhaB is secreted across the OM by FhaC and emerges as a hairpin on the eel surface. The N-terminal ACD domain of ACT binds to the FhaB MCD forming a stable complex. Engagement of FhaB with an unknown receptor and ACT with its receptor (CR3) causes a signal to be transduced that initiates regulated degradation of the periplasmic FhaB prodomain by DegP and CtpA. Prodomain degradation causes the C terminus of processed FhaB to exit the FhaC channel culminating in delivery of ACT to the CR3+ host cell.

ACT is a large (1,706 aa) bifunctional protein. Its N-terminal adenylate cyclase domain binds calmodulin and catalyzes the production of cAMP from ATP(26,27). Its C-terminal RTX hemolysin domain contains (from N terminus to C terminus) a hydrophobic pore-forming subdomain, a segment that is palmitoylated on lysine residues by the *cyaC* gene product (28,29), a calcium-binding RTX subdomain composed of 45 nonapeptide repeats, and a signal for export by the Type I secretion system formed by the products of the *cyaB*, *cyaD*, and *cyaE* genes(30,31). A DDE motif within the nonapeptide repeats mediates binding of ACT to the CD11b component of CR3 (CD11b/CD18, Mac1), a receptor present on macrophages, dendritic cells, and neutrophils(32).

ACT, via its AC domain, binds FhaB on the surface of *B. pertussis* and *B. bronchiseptica*, and this binding prevents FhaB-FhaB interactions that are critical for biofilm formation(7). Hence, *Bordetella* species form robust biofilms when growing in BvgAS-intermediate (BvgAS^i^) mode and produce FhaB but not ACT(8,33). (BvgAS is a two-component regulatory system that activates expression of all protein virulence factor-encoding genes. When it is partially active (BvgAS^i^ mode), it activates expression of genes with high affinity BvgA binding sites at their promoters, such as *fhaB*, but not those with low affinity BvgA binding sites, such as *cyaA*.) Strains lacking ACT also form biofilms under BvgAS^+^ mode conditions(7). Antibodies against the MCD of FhaB block ACT from binding to FhaB(7), suggesting that the AC domain binds specifically to the FhaB MCD, but details about the ACT-FhaB interaction are lacking.

We showed several years ago that *B. bronchiseptica* strains producing FhaB proteins lacking the ECT or PRR (at the C terminus of FhaB) are defective for persistence in the lower respiratory tract (LRT), despite adhering to host cells as effectively as wild-type bacteria(12). Because the PD is degraded in the periplasm and does not function as a stand-alone polypeptide, these data indicate that full-length FhaB plays an important role in defense against phagocytic cells in the LRT during infection. Given the importance of ACT in defense against phagocytic cells, and the fact that it binds to FhaB on the bacterial cell surface, we have speculated that FhaB may serve as a novel toxin delivery system that controls delivery of ACT specifically to phagocytic cells in the respiratory tract(13,23). According to this model (Fig. 1), newly secreted ACT preferentially binds FhaB (that is in complex with FhaC) on the bacterial surface. Binding of ACT to CR3 on phagocytes generates a signal that is propagated across the outer membrane, resulting in a conformational change in the FhaB PD such that DegP cleaves off the ECT, revealing a C terminus that is degraded by CtpA. Degradation of the PD, we hypothesize, allows the C terminus of FHA’ to slide through FhaC, which allows ACT to be released from FhaB and delivered to the phagocytic cell. Here, we present evidence that supports this model, that identifies ACT as a substrate for SphB1, and that shows that FhaB-bound ACT can be delivered to host cells.

## RESULTS

### ACT binds FhaB preferentially before being released from the surface of *B. bronchiseptica*

ACT interacts with FhaB on the surface of both *B. bronchiseptica* and *B. pertussis*(1,7). However, studies identifying these interactions did not examine the dynamics of ACT-FhaB binding. Our model predicts that ACT binds FhaB preferentially before being released into culture supernatants. To test this hypothesis, we sought to track the amount and localization of newly secreted ACT in wild-type and Δ*fhaB* bacteria. We degraded surface-associated, external ACT with Proteinase K (Prot K), washed the bacteria to remove the enzyme and supernatant polypeptides, sub-cultured bacteria in fresh medium, and compared the amount of ACT on the bacterial surface, in culture supernatants, and in whole cell lysates (WCL) via dot and western blot analyses.

To determine the appropriate concentration of Prot K for these experiments, we treated wild-type bacteria with 0-8 μg/mL Prot K for 30 minutes, washed bacteria thoroughly, and performed dot blots on intact bacteria using FhaB- and ACT-specific antibodies. 4 μg/mL Prot K was sufficient to degrade ACT to levels undetectable by dot blot while leaving FhaB largely intact due to its intrinsic resistance to proteolysis (Figure 2A). We used this concentration of Prot K for subsequent experiments.

**Figure 2.**
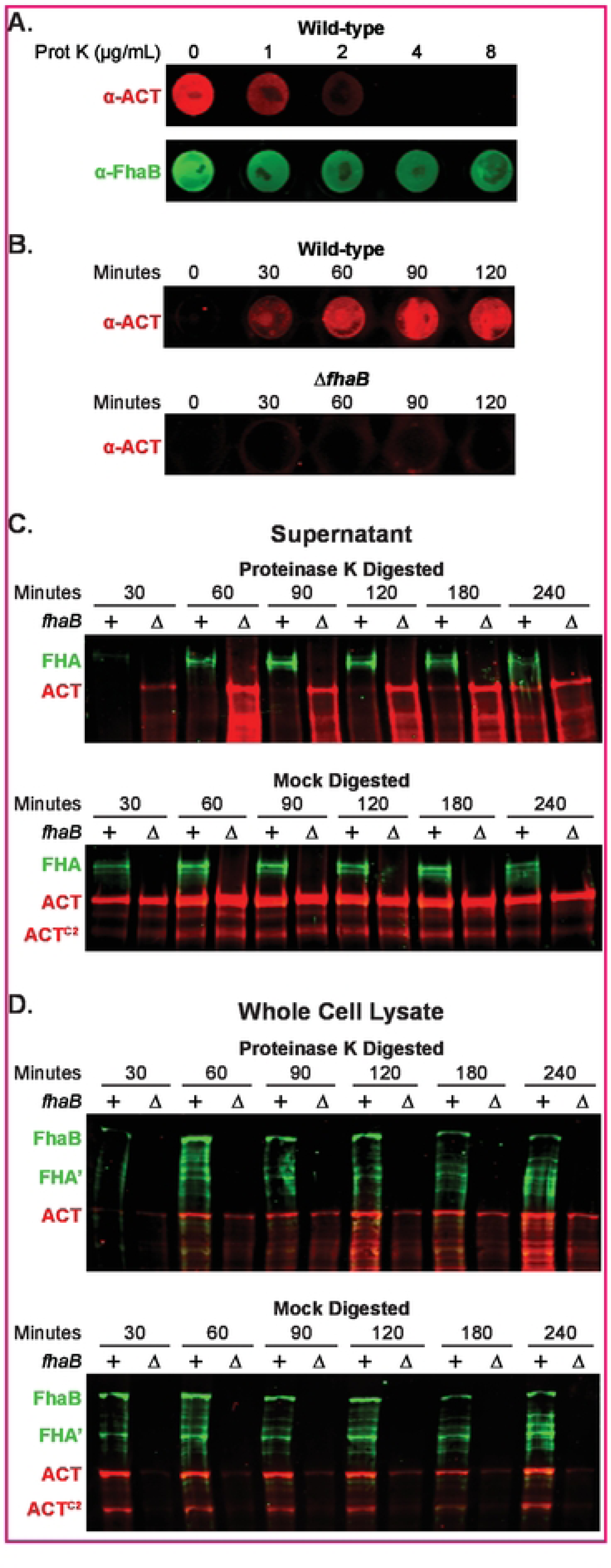
Newly secreted ACT binds FhaB on the cell surface prior to release. A. To degrade surface exposed proteins, wild-type bacteria were treated for 30 minutes with varying concentrations of Proteinase K (Pro! K) then washed. Surface ACT and FhaB were probed for via dot blot of intact cells using polyclonal a-ACT and a-FhaB antibodies. 8. Wild-type and*!lfhaB* strains were treated with 4 µg/ml of Pro! K for 30 minutes, washed, and cultured further in fresh media. Bacteria were sampled at the times listed to probe by dot blot for surface-associated ACT. C and D. Wt and *llfhaB* strains were treated with 4 µg/ml of Prot K for 30 minutes (or mock digested with solution lacking Prot K), washed, and subcultured for the times listed. Production, processing, and release of FhaB and of ACT were examined by western blot analyses of filtered supernatants (C) and of whole cell lysates (D), using polyclonal a-FhaB and monoclonal a-ACT antibodies that recognize the MCD and the AC domain C terminus, respectively.

We next examined the amount of time required for newly secreted ACT to accumulate at the bacterial surface to levels detectable by dot blot following treatment with Prot K. Intact bacteria collected at 0, 30-, 60-, 90-, and 120-minutes post-treatment were probed with ACT-specific polyclonal antibodies. ACT was detected on the surface of wild-type bacteria beginning at 30 minutes, and the amount increased for the following hour. By comparison, no ACT was detected on the surface of the Δ*fhaB* strain even after 120 minutes of post-treatment culturing, supporting previous findings that FhaB is required to retain ACT on the *B. bronchiseptica* surface (Fig. 2B). In culture supernatants of Δ*fhaB* bacteria (normalized to the OD_600_ of the culture), ACT was detectable by western blot at 30 minutes post-Prot K treatment and increased over time (Fig. 2C). By contrast, ACT was not appreciably detected in culture supernatants of wild-type bacteria until 180 minutes post-Prot K treatment, and did not match levels present in supernatants of Δ*fhaB* bacteria until 240 minutes post treatment (Fig 2C). Without Prot K digestion, amounts of ACT in culture supernatants of wild-type and Δ*fhaB* bacteria were equivalent at every time point.

In contrast to supernatants, cell-associated ACT was detected in WCL of wild-type and Δ*fhaB* bacteria at 60 minutes post-treatment with Prot K, and all timepoints after that (Fig. 2D). In all cases, the amount of ACT present in WCL of the Δ*fhaB* strain was less than that present in WCL of wild-type bacteria. The ACT detected in WCL of the Δ*fhaB* mutant was likely newly synthesized, cytoplasmic ACT as it was not detected by dot blot (Fig. 2A). WCL of mock-treated wild-type and Δ*fhaB* bacteria secreted similar amounts of ACT at all times tested. Collectively these data indicate that secreted ACT preferentially binds FhaB on the cell surface until ACT has bound all of the available FhaB, at which point excess toxin is released directly or in complex with FhaB into the medium.

### ACT inhibits FhaB processing

To determine if the presence of ACT affects processing of FhaB, we compared FhaB and ACT profiles in wild-type and ACT-deficient strains by western blot and dot blot analyses using anti-FhaB (green in Fig. 3) and anti-ACT (red in Fig. 3) antibodies. Because it has been reported that Ca^2+^ concentration in the medium influences ACT localization(34), we grew bacteria in either standard Stainer Scholte (SS) broth(35), which contains 0.18 mM Ca^2+^, or in SS broth containing 2.0 mM Ca^2+^. ACT was detected on the surface of wild-type, but not Δ*fhaB* or Δ*cyaA*, bacteria grown in both media (Fig. 3A), indicating that ACT remains associated with the bacterial surface in an FhaB-dependent manner regardless of Ca^2+^ concentration.

**Figure 3.**
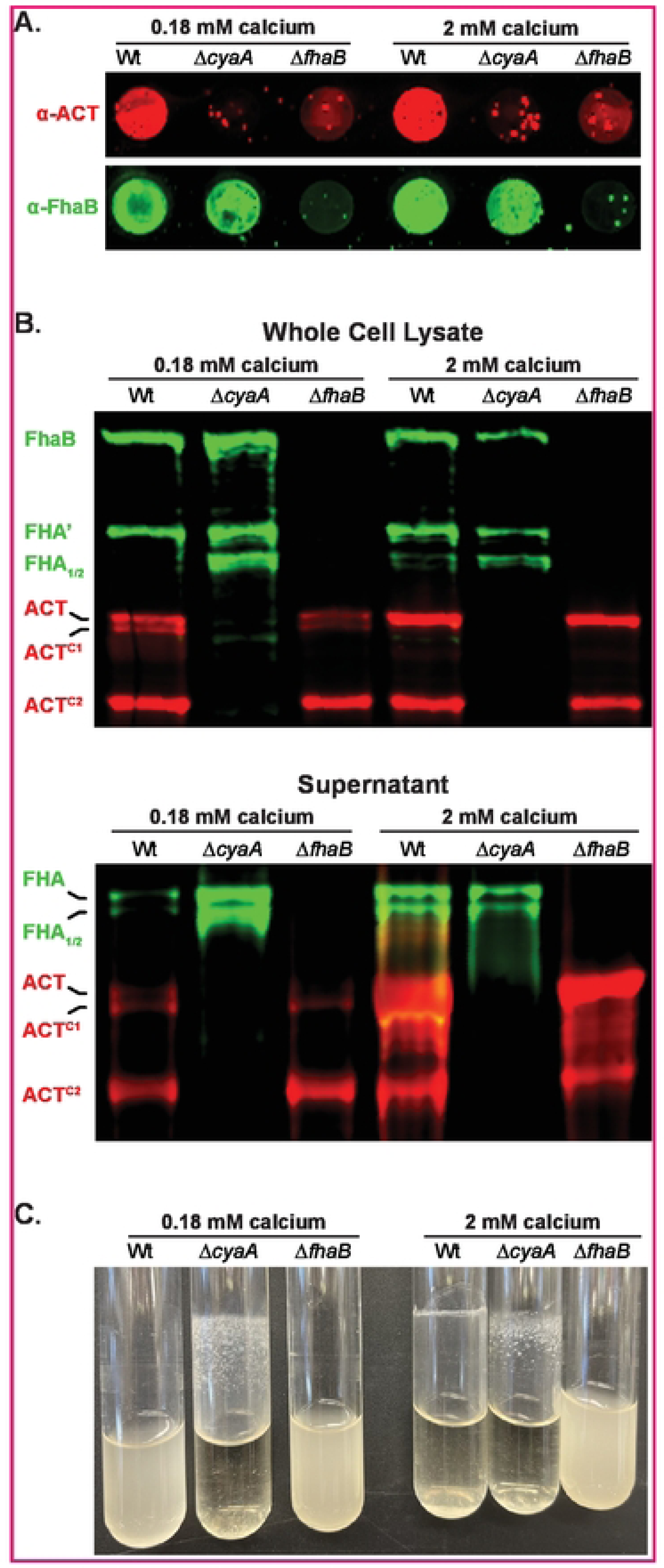
Calcium does not prevent association of FhaB and ACT on the bacterial surface of *B. bronchiseptica*. A. Dot blots of intact bacteria cultured for 20 hours in standard Stainer Scholle (SS) broth, which has 0.18 mM calcium, or SS supplemented with 1.8 mM additional calcium (bringing the total to 2 mM calcium). Blots were probed with polyclonal a-ACT and a-FhaB B. Western blot analyses of whole cell lysates and culture supernatants grown in conditioned mentioned in A. Blots were probed with polyclonal a-FhaB and 3D1 monoclonal a-ACT antibodies. Full length and multiple cleaved forms of ACT and FhaB are indicated. C. Culture tubes of wild-type, *acyaA,* and *6fhaB* bacteria cultured for 20 hours in standard Stainer Scholle (SS) broth or SS supplemented with additional calcium. The adherent band occurs at the liquid-air-glass interface while rotating tilled.

WCL of wild-type bacteria typically contain full-length FhaB and FHA’, the FhaB polypeptide remaining after DegP- and CtpA-mediated degradation of the prodomain ((23,24), Fig. 3B, and see Fig.1). In the absence of ACT (Δ*cyaA*), FHA_1_ and FHA_2_, the products of SphB1-dependent cleavage, were also present (Fig. 3B). Culture supernatants of wild-type bacteria typically contain FHA, FHA_1_, and FHA_2_, and all of these polypeptides were more abundant in culture supernatants of the Δ*cyaA* strain compared to wild-type bacteria when grown in medium containing 0.18 mM Ca^2+^. Consistent with Bumba *et al*.(34), there was substantially more ACT in culture supernatants of bacteria grown in 2.0 mM Ca^2+^ (despite there being similar levels in WCL). Our data indicate that supplementation with 2.0 mM Ca^2+^ results in greater secretion of ACT into the medium, but that a substantial amount of ACT remains associated with FhaB, and that association of ACT with FhaB, especially in 0.18 mM Ca^2+^, inhibits or blocks SphB1-dependent cleavage of FhaB. The fact that there is so much more FHA (which is formed only after DegP- and CtpA-dependent degradation of the prodomain) in culture supernatants of the Δ*cyaA* strain grown in standard SS indicates that the presence of ACT also affects SphB1-independent processing of FhaB.

Wild-type and Δ*fhaB B. bronchiseptica* grow planktonically in standard SS medium, while the Δ*cyaA* strain grows as a biofilm adherent to the walls of the culture tube (Fig. 3C). When grown in SS containing 2.0 mM Ca^2+^, wild-type bacteria also aggregated and formed a ring in the culture tubes, but in a way that was different from the biofilm formed by the Δ*cyaA* strain (Fig. 3C). These data indicate that the presence of 2.0 mM Ca^2+^ alters, but does not abrogate, the FhaB-ACT interaction. We hypothesize that 2.0 mM Ca^2+^ prevents ACT-ACT interactions, but does not completely abrogate FhaB-ACT interactions on the bacterial surface.

### ACT is cleaved in an SphB1-dependent manner

Although the primary amino acid sequence of ACT predicts it to be a 177 kD polypeptide, ACT typically runs with the mobility of a larger ∼200 kD protein by SDS-PAGE (ACT in Figs. 2 & 3). A substantial amount of a polypeptide running with an apparent MW of ∼175 kD (ACT^C2^) was also detected in the western blots shown in Figs. 2 & 3. To investigate the nature of those polypeptides, we probed WCL of wild-type *B. bronchiseptica* with a monoclonal antibody that recognizes an epitope near the C terminus of the AC domain (3D1)(36), a monoclonal antibody that recognizes an epitope within the RTX domain (9D4)(36), and a polyclonal antibody raised against the entire ACT protein (Poly). The most prominent polypeptides, ACT, ACT^C1^, and ACT^C2^, as well as a less abundant polypeptide running at about 140 kD (ACT^C4^), were recognized by all three antibodies (Fig. 4B), indicating that those polypeptides had lost aa at the N terminus of ACT. A polypeptide running at ∼170 (ACT^C3^) was recognized by the 9D4 monoclonal antibody and the polyclonal antibody, but not 3D1 (Fig. 4A), indicating ACT^C3^ contained the C terminus of ACT and not the AC domain. Based on preliminary experiments, we suspected that SphB1 may be involved in cleavage of ACT. Western blot analysis with the 3D1 antibody showed that ACT was the predominant or only form of ACT in WCL and supernatants of Δ*sphB1* bacteria (Fig. 4C), demonstrating that cleavage of ACT is indeed SphB1-dependent. These data represent the first demonstration that SphB1 cleaves a protein other than itself and FhaB(25).

**Figure 4.**
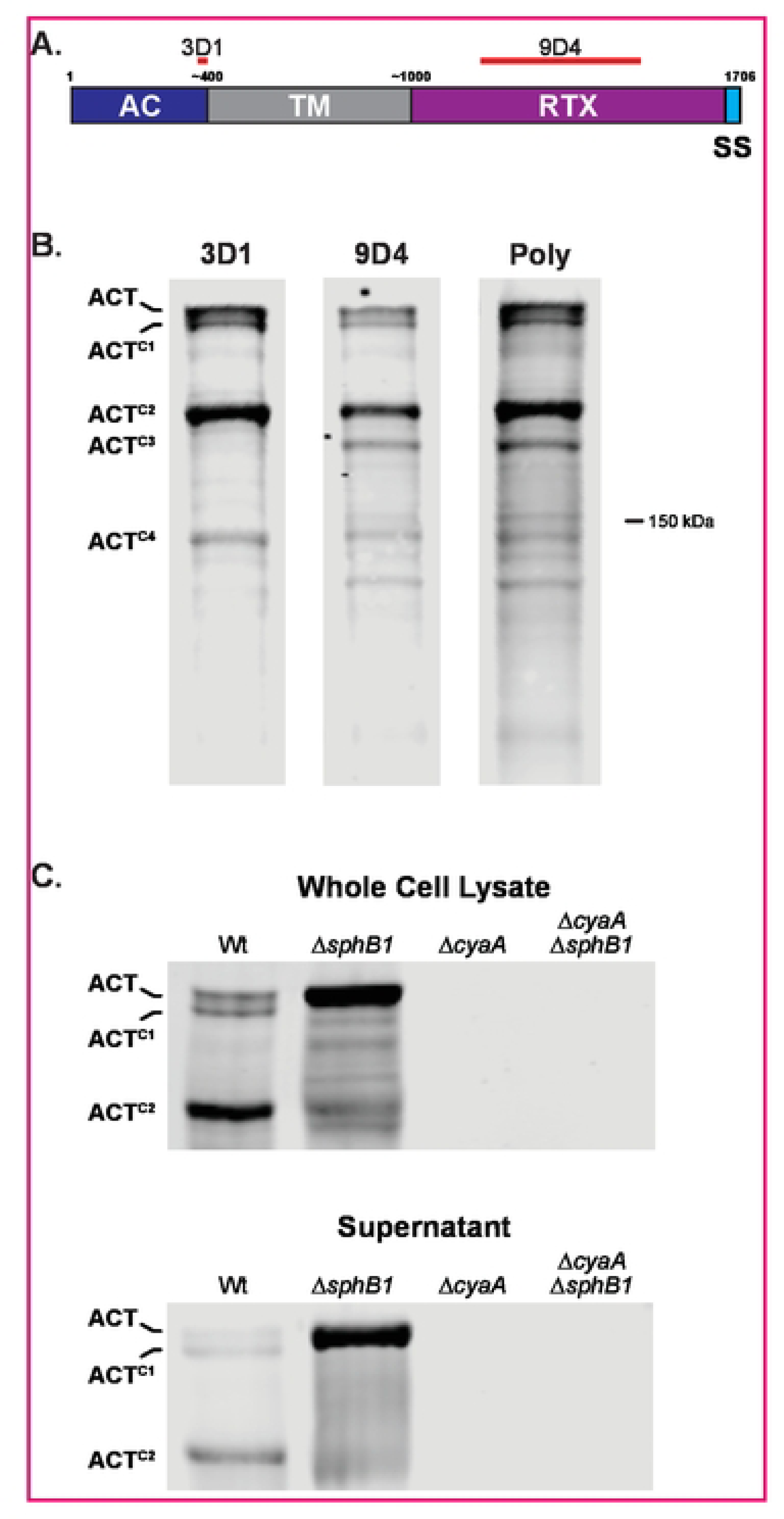
ACT is cleaved in an Sph81-dependent manner. A. Schematic of ACT, showing adenylate cyclase domain (AC), transmembrane domain (TM), repeats-in-toxin domain (RTX), and C-terminal signal sequence (SS). Estimated and actual amino acid markers are shown at top. Regions used to generate monoclonal antibodies 3D1 and 9D4 are indicated with red bars overlapping residues 373-399 and 1156-1489, respectively. B. Western blots of whole cell lysates from wild-type bacteria were probed with monoclonal antibodies 3D1 or 9D4 or with polyclonal antibody generated using whole protein (Poly). C. Western blot analyses of *B. bronchiseptica* strains lacking SphB1 and/or ACT. Blots are focused on three forms in both whole cell lysates and culture supernatants: uncleaved ACT and two cleaved forms ACTc^1^ and ACTc^2^ Blots were probed with 3D1 a-AC domain monoclonal antibody.

### SphB1-dependent cleavage of ACT is predicted to inactivate ACT

To characterize the SphB1-dependent ACT cleavage products further, we constructed strains producing ACT with an HA epitope at either the N or C terminus and examined protein profiles via western blotting with the 3D1 antibody (red in Fig. 5) and polyclonal rabbit anti-HA antibodies (green in Fig. 5). The N-terminal HA epitope was barely detected in the intact ACT polypeptide (and not detected in the smaller ACT polypeptides) (Fig. 5, and associated supplemental Fig), indicating that the N-terminal HA epitope was proteolytically removed from a majority of the ACT polypeptides. The C-terminal HA epitope was present on the full-length ACT_CT-HA_ protein, ACT^C1^, ACT^C2^, and ACT^C3^, indicating that SphB1-dependent cleavage to produce ACT^C1^ and ACT^C2^ occurs within the AC domain (Figure 4A). ACT^C3^ was not detected by the 3D1 antibody, indicating that cleavage to produce this polypeptide occurs C-terminal to the 3D1 recognition site (i.e., within the TM domain). Consistent with the western blot results, the C-terminal, but not N-terminal, HA epitope was detected on the cell surface by dot blot (Fig. 5B). These data indicate that SphB1-dependent cleavage of ACT occurs at three or more distinct sites within ACT.

**Figure 5.**
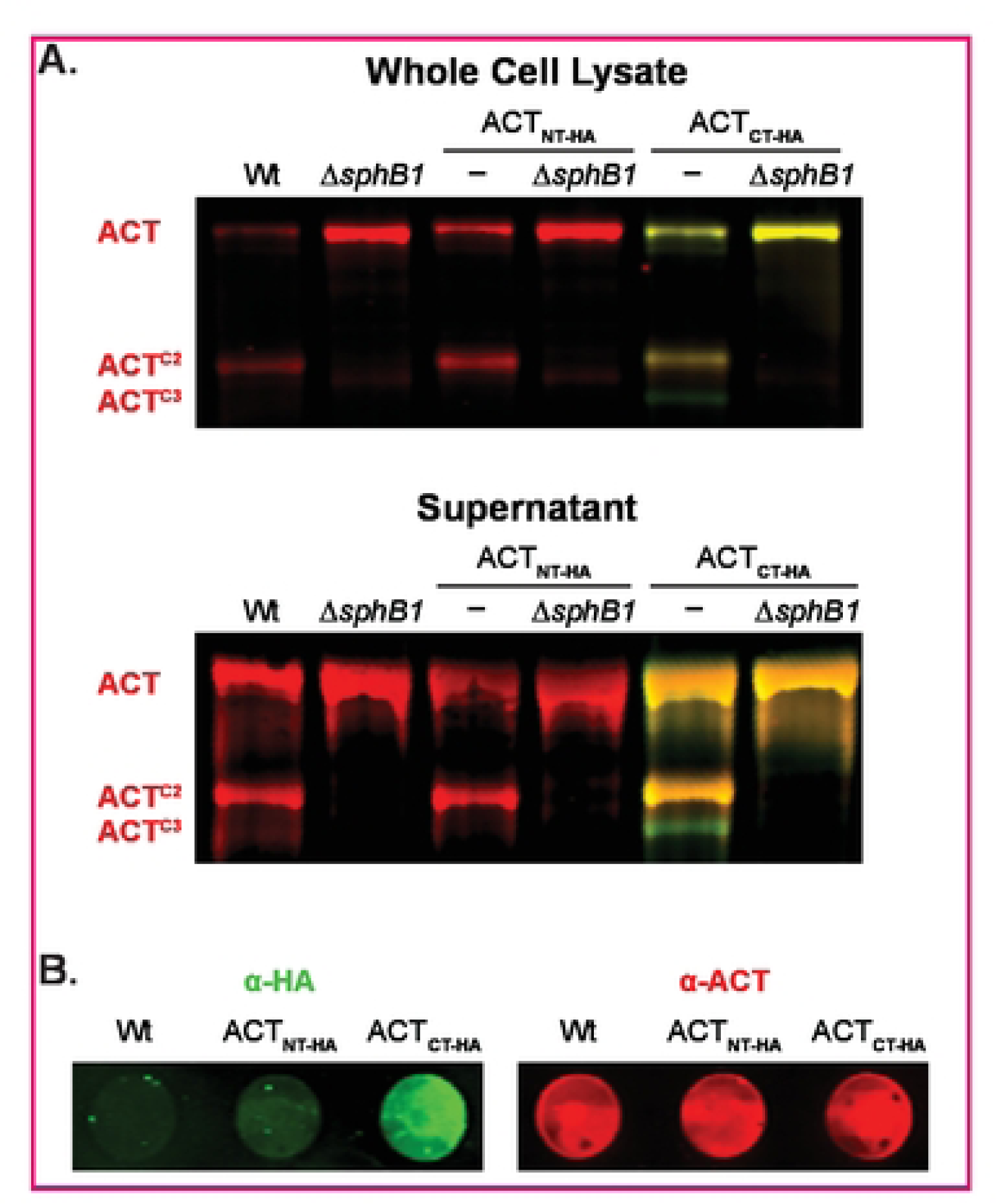
SphB1-dependent cleavage of ACT occurs at the N-terminus. A. Western blot analyses of HA-tagged ACT, with the tag at either the N or C terminus (ACTNT-HAand ACTcr-HA• respectively). Red bands indicate ACT polypeptides that possess residues 373-399 (the AC domain C terminus recognized by antibody 301, see Figure 4). Yellow bands indicate polypeptides that possess ACT residues 373-399 and also an HA tag. Green bands indicate polypeptides that lack residues 373-399 but possess an HA-tag. B. Dot blots of intact wild-type bacteria and strains that produce HA-tagged versions of ACT. Bacteria were seperately probed for HA (green) or ACT (red) using polyclonal antibodies.

ACT^C2^ was the most abundant ACT polypeptide in both supernatants and WCLs of overnight cultures of wild-type bacteria. We used Edman sequencing to determine that the ACT^C2^ N terminus is TRLGQLKEY, which matches the aa sequence spanning T326-Y334 within the ACT N-terminal ACD. This indicates that the SphB1-dependent cleavage event that generates ACT^C2^ occurs between L325 and T326 (dashed line in Figure 6B), and that ACT^C2^ contains the anti-3D1 epitope. We used allelic exchange to construct three strains producing ACT with aa substitutions in and around the identified cleavage site (changed residues shown in red in Fig. 6B). The SphB1-dependent ACT^C2^ polypeptide was not detected in WCLs or supernatants from *B. bronchiseptica* producing the altered forms of ACT (Figure 6C,D), indicating that the aa substitutions abrogated SphB1-dependent cleavage of ACT at this site. As expected, ACT^C1^ was present in WCLs of all of the mutants, and was absent in the GVIDVE mutant in which *sphB1* was mutated (Fig. 6C). Dot blots of these strains using ACT- and FhaB-specific antibodies showed similar amounts of ACT^GVIDVE^, ACT^M324D/L325P^(DP), and ACT^M324Y/L325F^ (YF) on the surface of the mutant strains as ACT on wild-type bacteria, indicating that the aa substitutions did not disrupt the association between the ACD and FhaB on the bacterial surface (Supp associated with Fig 6).

**Figure 6.**
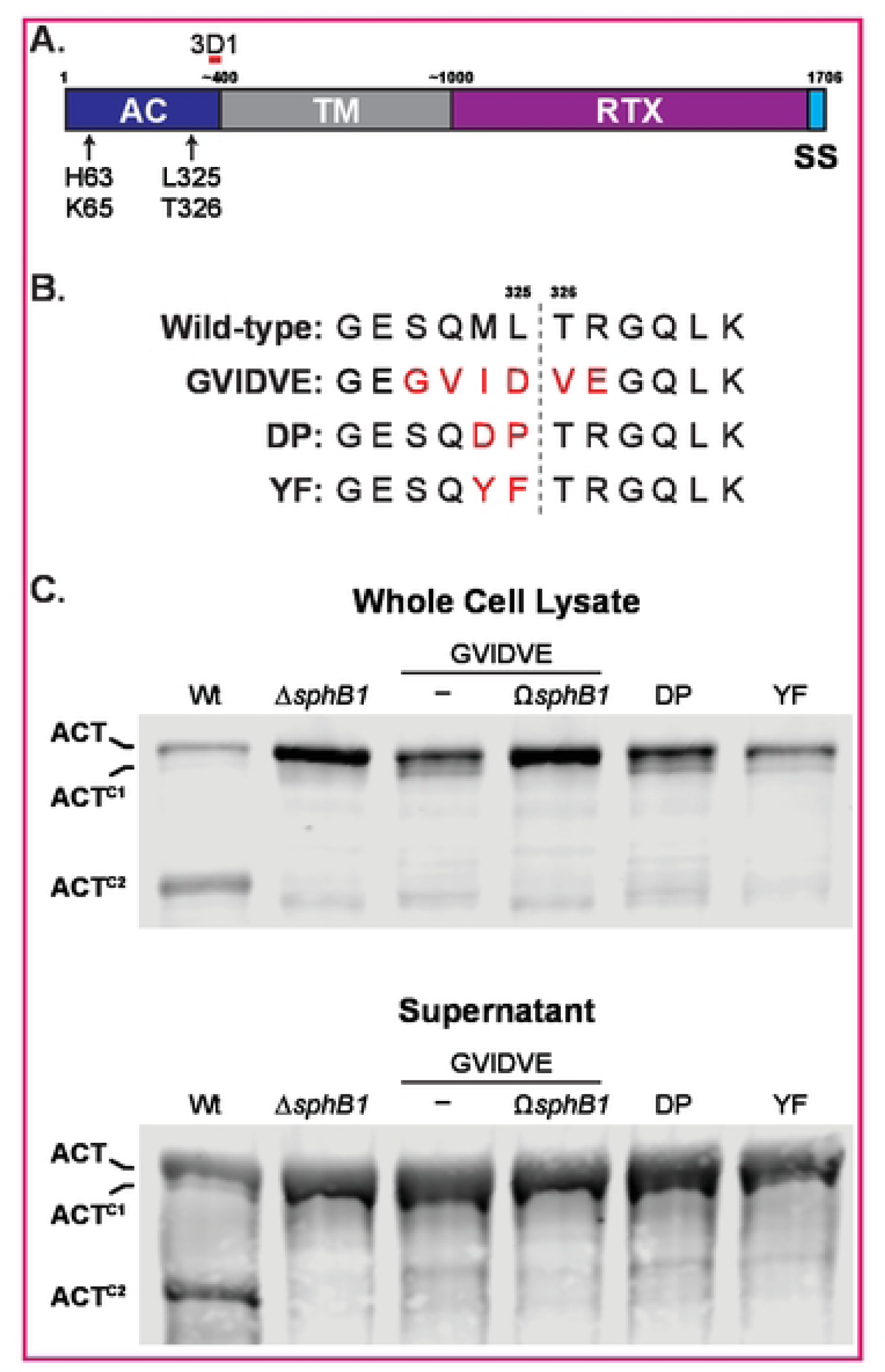
Identification of the Sph81 cleavage site on ACT. A. Schematic of ACT, showing adenylate cyclase domain (AC), transmembrane domain (TM), repeats-in-toxin domain (RTX), and C-terminal signal sequence (SS). Estimated and actual amino acid markers are shown at top. AC catalytic residues histidine 63 and lysine 65 are indicated, as well as the predicted SphB1-dependent cleavage site between leucine 325 and threonine 326 that yields ACTC2. Monoclonal antibody 3D1 recognition region is indicated by a short red bar overlapping residues 373-399. B. Sequence of the predicted SphB1-dependent cleavage site within the AC domain compared between wild-type ACT and three amino acid substitution mutants. Substitutions are highlighted in red, and the cleavage location between L325 and T326 is marked by a gray dotted line. C. Western blot analyses of whole cell lysates and culture supernatants of strains with ACT modifications as shown in B. Also included are wild-type, *6sphB1,* and GVIDVE with an insertional disruption of *sphB1* (GVIDVE *OsphB1).* Blots were probed with 3D1 antibody, and the three primary ACT forms are marked at left.

### SphB1-dependent cleavage of ACT does not require surface retention of ACT by FhaB

To determine if SphB1-dependent cleavage of ACT requires ACT’s interaction with FhaB, we compared ACT protein profiles in wild-type and Δ*fhaB* strains. For these experiments, we grew bacteria in media containing 50 mM MgSO_4_, a condition in which BvgAS, the two-component regulatory system that activates expression of *fhaB* and *cyaA,* is inactive, then switched the culture to BvgAS-activating conditions and collected samples at four- and 16-hours post-shift. At four hours post-shift, ACT, and ACT^C2^ were present in WCL of wild-type and Δ*fhaB* strains (Fig. 7A), indicating that SphB1-dependent cleavage to form ACT^C2^ occurs even in the absence of FhaB. At 16 hours post-shift, ACT^C1^ was also present, suggesting cleavage to form ACT^C1^ also occurs in the absence of FhaB, but is less efficient than cleavage to form ACT^C2^. These data indicate that SphB1-dependent cleavage of ACT does not require retention of ACT on the surface by FhaB.

**Figure 7.**
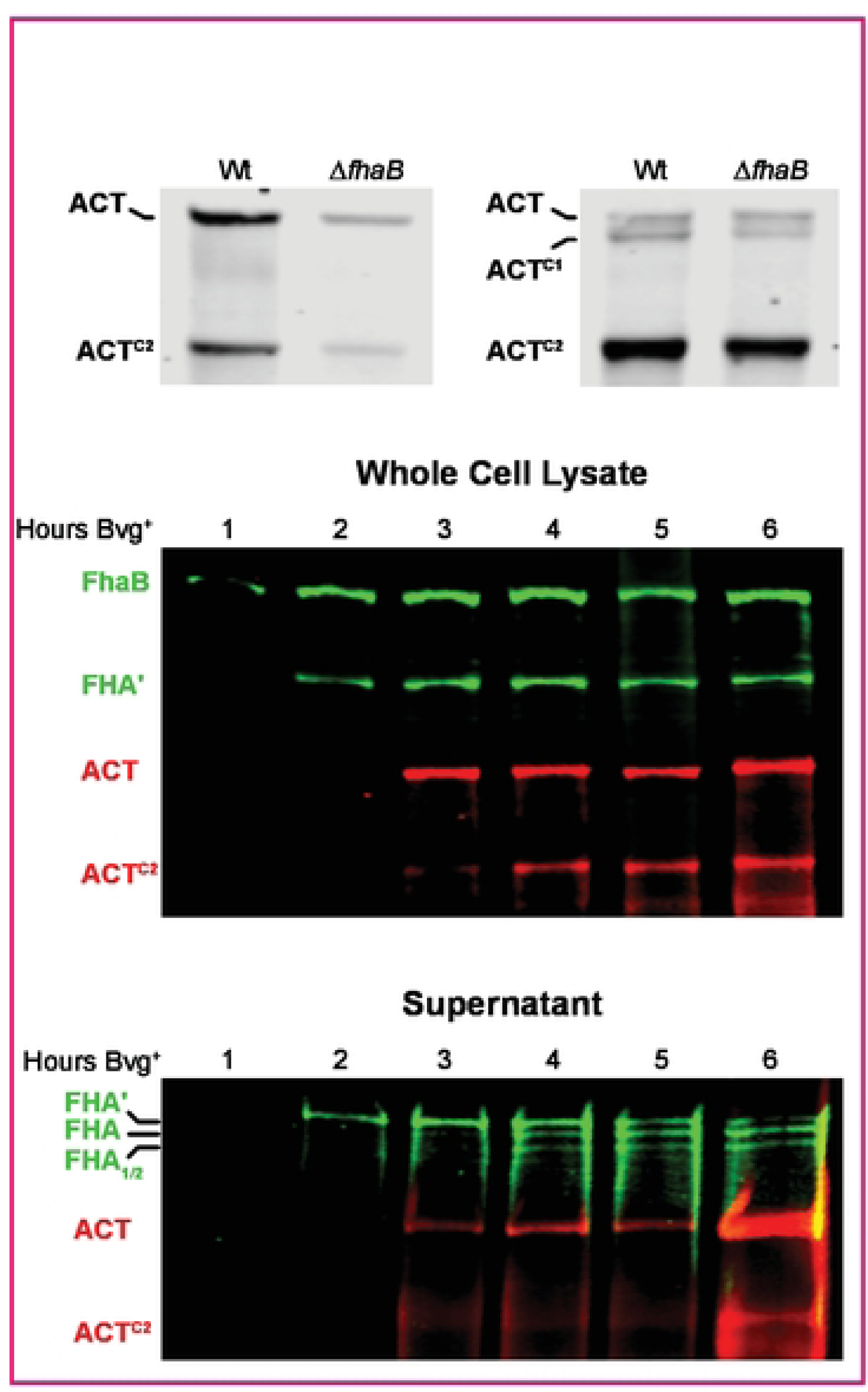
Production and degradation of FhaB and ACT overtime. A. ACT degradation in whole cell lysates of wild-type bacteria that were grown for 4 hours or 16 hours in BvgAS inducing media. ACT was probed via western blot with a-ACT monoclonal antibody. B. Production, degradation, and release of FhaB and ACT through 6 hours post-induction. Wild-type bacteria were subcultured in BvgAS inducing media (SS media) following overnight growth in non-inducing media (SSmedia with 50mM MgS0_4_), and whole cell lysates and culture supematants were probed via western blot analyses with a-FhaB polyclonal and a-ACT monoclonal antibodies. Full length proteins and the major degraded polypeptides are indicated.

To investigate maturation of FhaB and ACT in more detail, we monitored FhaB and ACT protein profiles every hour for the first six hours post-shift from BvgAS-inactivating to BvgAS-activating conditions. In WCL, only full-length FhaB was detected at one-hour post-shift (Fig. 7B). FHA’, resulting from degradation of the prodomain by DegP and CtpA, was detectable at 2 hours post-shift, and ACT was first detected at 3 hours post-shift. ACT^C2^ was detected at 4 hours post-shift. In culture supernatants, SphB1-dependent FHA and FHA_1/2_ were detected at about 3 – 4 hours post-shift, while ACT^C2^ was not detected until about 6 hours post-shift. These data indicate a sequence of events in which FhaB is produced, then processed by DegP and CtpA and then SphB1, and then ACT is produced and then cleaved by SphB1.

### *B. bronchiseptica* delivers FhaB-bound-ACT to CR3+ mammalian cells

Our data are consistent with a model (Fig. 1 and Fig. 10) in which ACT binds to FhaB immediately upon secretion to the cell surface, and that in response to ACT binding its receptor (CR3), FhaB prodomain degradation allows ACT to be delivered to the phagocytic cell, wherein the AC domain catalyzes the production of cAMP from ATP. To investigate intoxication of host cells by *B. bronchiseptica*, we infected J774A.1 murine macrophage-like (CR3+) cells, Chinese hamster ovary cells (CHO, CR3-), and CHO cells engineered to produce human CD18 and CD11b (CHO CR3+(32)) with wild-type and Δ*cyaA B. bronchiseptica* strains and measured cAMP levels using a competitive ELISA. As expected, wild-type *B. bronchiseptica* efficiently intoxicated CR3+ cells (J774A.1 and CHO+CR3) but not the CR3– CHO cells (Fig. 8) (or A549 human epithelial CR3– cells, not shown), indicating that the CR3 receptor greatly enhances entry of *B. bronchiseptica* ACT into host cells, similar to what has been shown for ACT from *B. pertussis*(32,37). As expected, no increase in cAMP occurred with the Δ*cyaA* strain.

**Fig 8.**
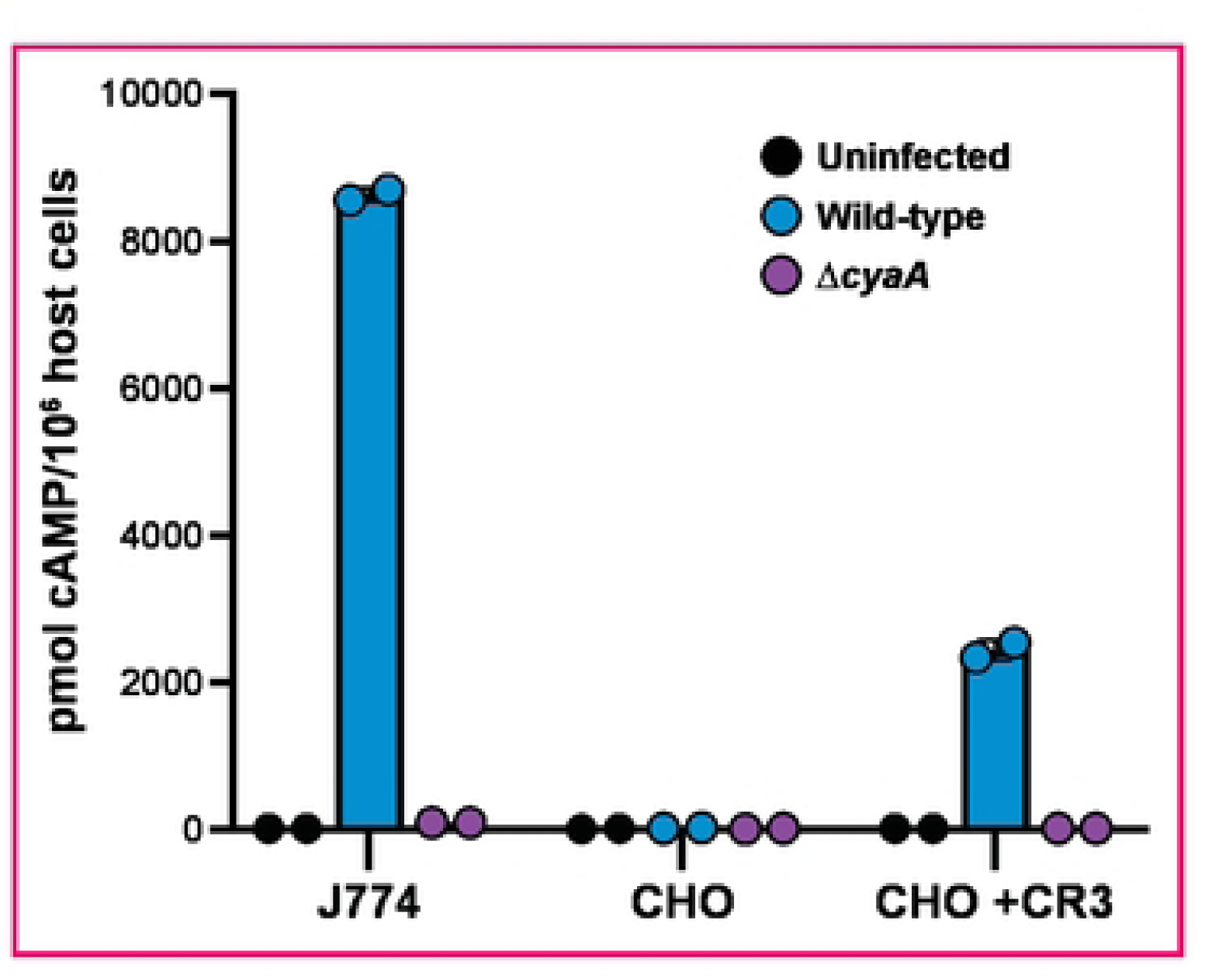
*B. bronchiseptica* intoxicates CR3-positive cells with ACT. J774A.1 macrophage cells (J774) and CHO-K1 Chinese hamster ovary cells containing integrated empty vector (CHO) or vector with DNA that encodes complement recepter 3 (CHO +CR3) were infected with wild-type and *6cyaA* bacteria for 30 minutes at an MOI of 100. cAMP amounts were determined by direct ELISA.

**Figure 9.**
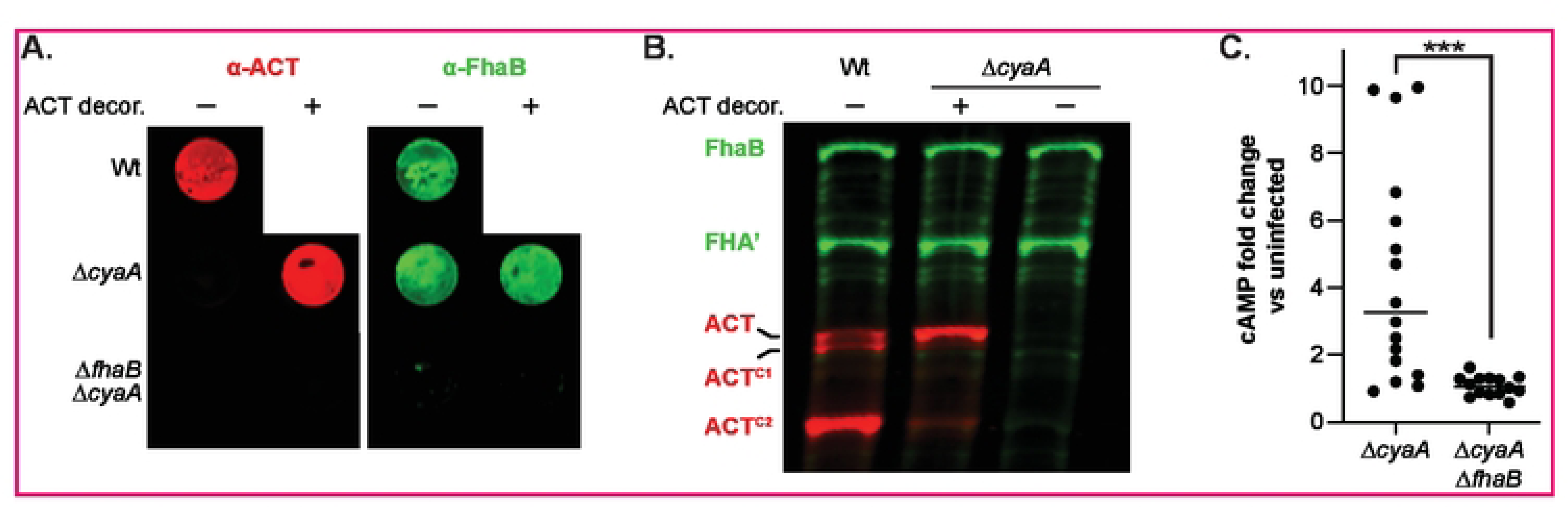
ACT bound to FhaB on the bacterial surface can be delivered to J774A.1 macrophage-like cells. A. Thesurfaces of ACT-null bacteria *(6cyaA* and*6cyaA6fha8* strains) were decorated with ACT released from *6fha8 6sph81* bacteria by suspending the ACT-null bacteria in sterile-filtered, ACT-containing culture supernatant and then washing in PBS to remove non-adhered ACT External, surface-bound ACT and FhaB were examined via dot blot of intact decorated and undecorated cells, probing with polyclonal a-ACT and a-FhaB antibodies. B. Western blot of Wild-type, decorated 6cyaA, and undecorated 6cyaA bacteria. FhaB, ACT, and their cleaved forms were detected using polyclonal a-FhaB and monoclonal a-ACT antibodies. C. J774.1 macrophages were infected with decorated bacteria for 15 minutes at an MOI of 100. Levels of cAMP were determined by ELISA and shown as fold change vs uninfected macrophages. The groups were compared using a student’s unpaired T-test (••• = p <0.001).

**Figure 10.**
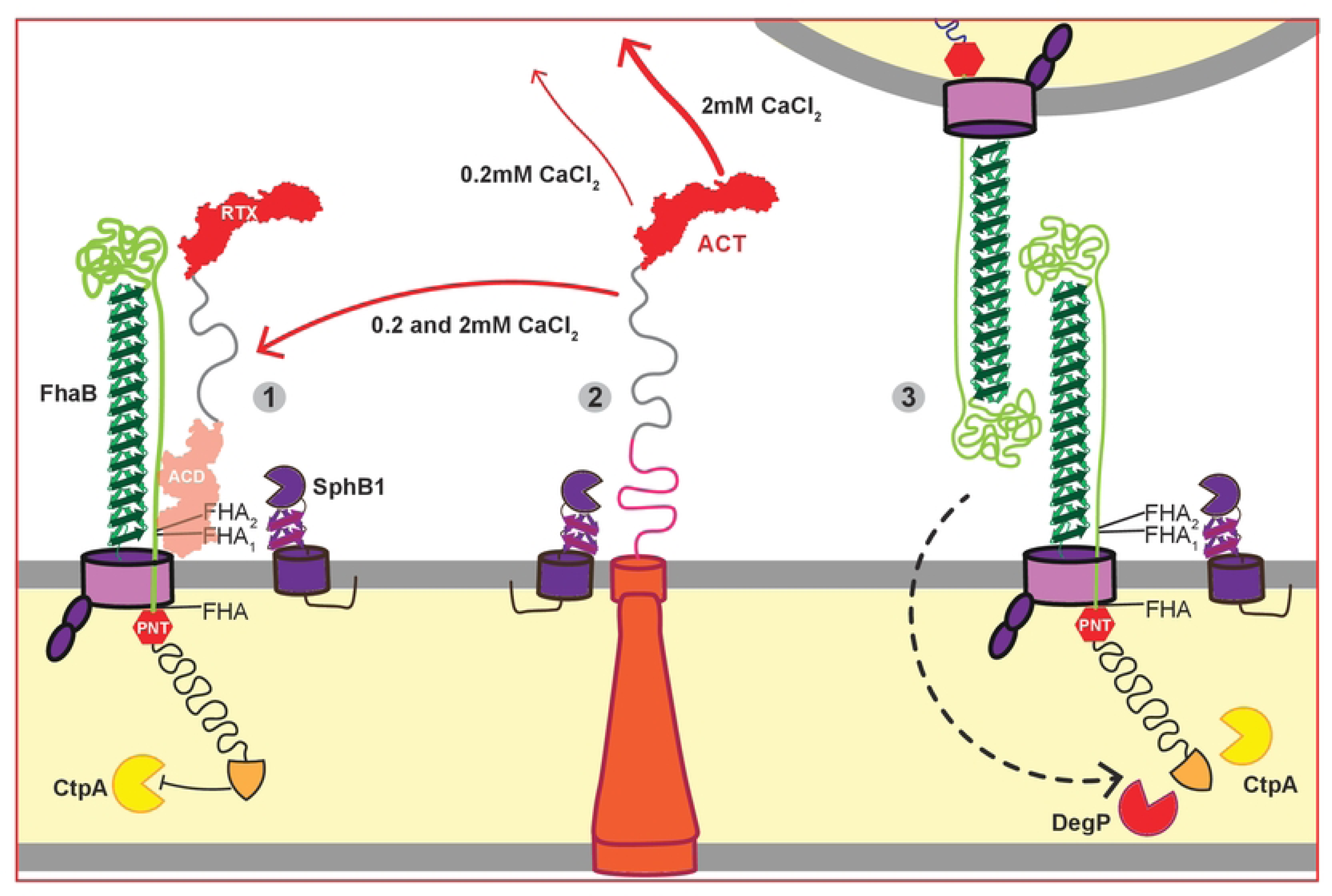
Model of the interactions between FhaB, ACT, and Sph81 on the surface of *Bordetella bronchiseptica*. 1. The N-terminal ACD of ACT binds the extracellular MCD of FhaB on the surface of wild-type bacteria blocking SphB1-dependent cleavage of FhaB at the FHA,_12_cleavage sites.
2. The rate of ACT secretion through the CyaBDE T1SS is higher when the bacteria are grown in SS broth with 2 mM than 0.2 mM CaCl_2_. In the absence of FhaB ACT undergoes Sph81-dependent cleavage at multiple sites within the ACD and TM domains on the cell surface.
3. In the absence of ACT, FhaB molecules on adjacent bacteria interact, resulting in bacterial aggregation and biofilm formation. ACT-deficient strains exhibit increased SphB1-dependent cleavage of FhaB to form FHA_112_ as well as increased DegP and CtpA processing of the FhaB prodomain.

It has been reported that FhaB-bound ACT is not delivered to host cells, i.e., that FhaB merely mediates adherence and it is newly secreted ACT that intoxicates the cell(38). To test directly the hypothesis that FhaB-bound ACT can be delivered to host cells, we incubated Δ*cyaA* strains with supernatants collected from cultures of the Δ*fhaB* Δ*sphB1* strain, which secretes large amounts of full-length extracellular ACT, such that the only ACT present on the bacteria would be that bound to FhaB on the cell surface (Figure 9). We confirmed the ACT ‘decoration’ of the Δ*cyaA* strain by examining ACT levels in WCLs and on the bacterial surface after thoroughly washing the decorated bacteria (Figure 9A). The Δ*cyaA* Δ*fhaB* strain did not retain ACT, indicating that FhaB is required for ACT decoration (Figure 9A). cAMP levels in J774A.1 cells incubated with the ACT-decorated Δ*cyaA* strain were significantly greater than J774A.1 cells incubated with the Δ*cyaA* Δ*fhaB* strain (Figure 9C). These data indicate that ACT bound to FhaB can be delivered to CR3+ host cells.

## DISCUSSION

Demonstration that ACT binds to FhaB on the surface of *B. pertussis* and *B. bronchiseptica* (1), together with the revelation that full-length FhaB plays an important role during infection(12), and that the FhaB prodomain is degraded in a regulated manner (23,24), prompted us to develop a model in which the binding of ACT to its receptor on a phagocytic cell triggers degradation of the FhaB prodomain and delivery of ACT to the host cell (Fig. 1). In this study, we investigated various aspects of the model.

Although previous studies showed that the N-terminal ACD of ACT binds to the MCD of FhaB in both *B. pertussis* and *B. bronchiseptica*(7), and that the binding is strong enough to require molar amounts of urea to remove ACT from FhaB(39–42), the nature of the interaction was not explored further. We showed here that ACT binds to FhaB preferentially on the bacterial surface before any ACT is released into the extracellular milieu (Fig. 2). Preferential and strong binding between ACT and FhaB suggests that the interaction between these proteins is important for the function of one, the other, or both during infection. Our data suggest that for FhaB, the interaction is primarily important for defense against clearance of the bacteria from the lower respiratory tract (LRT) by phagocytic cells, because FhaB mutants that result in hyper-degradation of the prodomain (and hence increased release of ACT) are able to adhere to host cells *in vitro* and *in vivo*, but are severely defective for persistence in the LRT(12). For ACT, we hypothesize that the interaction directs delivery of ACT to phagocytic cells, and not to epithelial cells or, wastefully, into the extracellular environment.

Although the interaction interfaces are not known, binding of the N-terminal ACD of ACT to the FhaB MCD supports the hypothesis that ACT sterically blocks SphB1-dependent cleavage of FhaB. Our data, showing increased amounts of FHA_1/2_ in WCL of the Δ*cyaA* strain compared to WCL of wild-type bacteria (Fig. 3B, WCL) also supports this hypothesis. Previous studies indicate that the ∼260 kD polypeptide in WCL of wild-type bacteria is FHA’ (the result of DegP- and CtpA-dependent processing) and not FHA (the result of SphB1-dependent cleavage at its primary site on FhaB)(24). Because they are so close in size, we do not know if the ∼260 kD polypeptide in WCL of the Δ*cyaA* strain is FHA’ or FHA. However, FHA is more readily released from the bacterial surface than FHA’(24), so, given the abundance of FhaB-derived polypeptides in the supernatants of the Δ*cyaA* strain (Fig. 3B, supernatants), it is likely that the larger band is composed of mostly, if not entirely, FHA. Because the formation of FHA requires DegP- and CtpA-dependent degradation of the prodomain (allowing the primary SphB1-dependent cleavage site on FhaB to move through FhaC and become exposed on the cell surface), these data suggest that the presence of ACT also influences DegP- and CtpA-dependent degradation of the prodomain. These data support the hypothesis that when ACT is bound to FhaB, the FhaB prodomain is in a stable conformation that is resistant to DegP-dependent processing. In the absence of ACT, or, we propose, when ACT binds CR3, the conformation of the prodomain changes such that DegP cleaves near the ECT, exposing a C-terminus that is susceptible to processive degradation by CtpA. These data, therefore, provide additional support for the model proposed in Fig. 1.

Our previous unpublished observations suggested that ACT may be cleaved in a SphB1-dependent manner. In this study, we showed that ACT is indeed cleaved at multiple sites in wild-type, but not Δ*sphB1*, bacteria, and we determined that the primary cleavage site is between L325 and T326. Prior analysis of the catalytic function of ACT indicates that cleavage at this location converts active ACT into two polypeptides, each incapable of converting ATP to cAMP. Efficient ACT catalytic activity requires both formation of the active site as well as binding to mammalian calmodulin (CaM). ACT digested with trypsin yields two inactive polypeptides: T25 – composed of aa1-235/237 including residues required for catalysis– and T18, which includes aa236/238-400(43). T18 and T25 do not readily associate, but can form a functional heterodimer in the presence of either CaM that binds the junction spanning the cleavage site(43,44), or if fused to interacting proteins – the basis of the first bacterial two-hybrid system(45). SphB1-dependent cleavage between L325 and T326 fully decouples the active site residues from the CaM binding domain and an ACT fragment spanning M324-R399 was unable to reconstitute adenylyl cyclase activity of an N-terminal ACT fragment(44). It is possible that ACT^C2^, while catalytically inert, can form pores in eukaryotic cells. However, the contribution of ACT-dependent pore formation to *Bordetella* pathogenesis has yet to be determined. Like Δ*cyaA* strains, *B. bronchiseptica* strains producing a catalytically inactive form of ACT are rapidly cleared from the murine lower respiratory tract(10), indicating that catalytic activity is the primary function of ACT during infection. Therefore, we interpret the finding that a majority of ACT is converted to catalytically inactive ACT^C2^ in overnight cultures of *B. bronchiseptica* to indicate that SphB1-dependent cleavage of ACT is the result of a degradative process rather than a maturation event, i.e., accumulation of the SphB1-dependent ACT^C2^ polypeptide on the cell surface may be an artifact of growing *B. bronchiseptica* for an extended time *in vitro*. Our previous work led to a similar conclusion for SphB1-dependent cleavage of FhaB(24), and hence we propose that SphB1 may function primarily as a ‘clean-up’ protease that removes damaged or ‘spent’ proteins from the bacterial surface.

Although the presence of ACT had a substantial effect on SphB1-dependent degradation of FhaB (especially when the bacteria were cultured in standard SS medium), the presence of FhaB had little to no effect on SphB1-dependent cleavage of ACT, indicating that FhaB does not block SphB1 from ACT and also that surface retention of ACT is not necessary for SphB1-dependent cleavage of ACT. Growth in medium containing 2 mM Ca^2+^ did, however, result in the release of dramatically more full-length ACT into the culture supernatant compared with growth in standard SS. These data are consistent with Ca^2+^ facilitating folding and secretion of the RTX domain of ACT(34), and suggest that in standard SS, insufficient Ca^2+^ slows the folding and secretion of ACT such that it is more accessible and more readily cleaved by SphB1.

Although the amount of ACT retained on the cell surface of bacteria grown in standard SS or SS containing 2 mM Ca^2+^ was similar (indicated by dot blot analysis), bacteria grown in SS containing 2 mM Ca^2+^ aggregated and formed a biofilm-like ring at the air-liquid interface of the test tube, suggesting increased FhaB-FhaB interactions and increased FhaB-mediated adherence to glass. The amount of biofilm and aggregation in wild-type bacteria grown in SS containing 2 mM Ca^2+^ was, however, less than the amount of biofilm and aggregation that occurred in cultures of Δ*cyaA* bacteria grown in either standard SS or SS containing 2 mM Ca^2+^. Together with results from Bumba et al.(34), our data suggest that although 2 mM Ca^2+^ facilitates folding, secretion, and release of ACT from the bacterial surface, it does not prevent a substantial amount of ACT from binding to FhaB on the cell surface.

A central component to our model is the hypothesis that *Bordetella* can deliver FhaB-associated ACT to CR3+ eukaryotic cells. Gray *et al*. investigated this hypothesis using *B. pertussis* and concluded that it is newly secreted, not FhaB-bound, ACT that is delivered to phagocytic cells(38). We used a different approach, decorating the surface of Δ*cyaA B. bronchiseptica* with ACT from supernatants of a Δ*fhaB* Δ*sphB1* donor strain and found that these bacteria could cause increased cAMP in J774A.1 macrophage-like cells. Because the only ACT present on these bacteria was that which was bound to the surface by FhaB, our data show definitively that FhaB can deliver ACT to host cells. However, intoxication of host cells by surface-associated (FhaB-bound) ACT or by ACT that has been secreted by its T1SS and avoided FhaB are not mutually exclusive. One possibility is that ACT bound to FhaB serves as a reservoir of active toxin that allows rapid inactivation of patrolling phagocytic cells, and that after delivery of the initial surface-associated ACT, the bacteria then continue to deliver more ACT to the host cell either via FhaB or directly from the T1SS.

A major challenge of all microbial pathogenesis research is reconciling results obtained from *in vitro* studies with the events that occur during infection. Our analyses have revealed interactions between FhaB, ACT, and SphB1, and are consistent with the model we have proposed in Fig. 1. Our model is also consistent with estimations by others that binding between FhaB and ACT is stronger (K_d_ of 6 x10^−7^)(7) than binding between ACT and CR3 (K_d_ of (1.2 x10^−3^)(32), which would mean that ACT could not be released from FhaB without a change in the conformation of one, or both, proteins (which, our model predicts, results from regulated degradation of the FhaB prodomain). The fact that *B. bronchiseptica* mutants that produce FhaB proteins that are hyperprocessed *in vitro* are severely defective for persistence *in vivo*(12) also supports the model. However, for at least two other TPS systems, the C-terminus of the TpsA protein is delivered to target cells (other bacteria, in the case of CdiA from *Escherichia coli*(46), host cells in the case of IbpA from *Histophilus somni*(47), and potentially both bacterial and host cells for CdiA from *Pseudomonas aeruginosa*(48)), and hence the possibility that binding to host cells triggers translocation of the FhaB prodomain through FhaC and delivery of the C-terminus to a host cell cannot, at this time, be ruled out.

## MATERIALS and METHODS

### Culture media and conditions

*Bordetella bronchiseptica* strains were streaked from −80°C stocks onto Bordet-Gengou agar (BD Biosciences) supplemented with 6% defibrinated sheep blood (HemoStat Laboratories) and grown at 37°C for 2-3 days. For broth cultures, colonies were picked from these plates and cultured overnight in Stainer-Scholte broth (SS). For experiments that monitored production and processing of nascent FhaB and ACT across time (such as Supp Assoc w/ Fig 6.), bacteria were instead first grown overnight in BvgAS non-inducing media (SS supplemented with 50 mM MgSO_4_) to prevent production of Bvg-induced virulence factors then washed with Dulbecco’s phosphate-buffered saline (PBS; Thermo Fischer) and subcultured in fresh BvgAS inducing media (standard SS) for times listed. *Escherichia coli* strains were grown at 37°C in lysogeny broth or on lysogeny broth agar. Where appropriate, media was supplemented with streptomycin (20 μg/mL), kanamycin (50 μg/mL), ampicillin (100 μg/mL), MgSO_4_ (50mM), and CaCl2 (1.8 mM).

### Bacterial strains and mutant creation

Bacterial strains and plasmids are listed in detail in Supplemental Table 1. *Escherichia coli* strains were used to amplify vectors (DH5α) and to transform (RHO3) *B. bronchiseptica*. In-frame deletions in *Bordetella bronchiseptica* were created via allelic exchange using derivatives of the pSS4245 plasmid, gene disruptions were created via insertion of the pEG7 plasmid within a coding region, and HA tags were added by allelic exchange. Mutations were confirmed by PCR and/or sequencing. As published in the past, we use three strains interchangeably as our wild-type: RB50, recovered from a naturally-infected rabbit; RBX11, an RB50 derivative in which *fhaB* is more genetically tractable because the strain lacks *fhaS*, a gene highly homologous to *fhaB* but that plays no role in virulence (Julio & Cotter, 2005); and RBX11 HA-PRR, an RBX11 derivative that produces FhaB with an HA-tag (YPYDVPDYA) inserted within the prodomain at a location that does not alter FhaB processing in any way that we have examined. Similarly, we often engineered specific mutations across more than one of these lineages. For ease of reading and because we have seen no lineage-dependent differences in regard to FhaB or ACT, we have not specified which lineage was used for specific experiments. Please note that FhaS does appear in culture supernatants when FhaB is absent (RB50 Δ*fhaB*).

### Immunoblots

To examine cell-associated proteins by western blot, whole cell lysates were prepared by boiling pelleted cells in Laemmli buffer. To examine released proteins, culture supernatants were filtered through 0.2μm filters, and proteins were precipitated using 10% trichloroacetic acid then rinsed with acetone, resuspended in 1M Tris-HCl pH 8.8 and Laemmli buffer mixture, and boiled. Proteins were separated by SDS-PAGE using 4% or 5% polyacrylamide gel and transferred to nitrocellulose membranes (GE Healthcare). Membranes were probed with mouse monoclonal antibody generated against HA-tag peptide (YPYDVPDYA; BioLegend), rabbit polyclonal antibody generated against the entire FhaB mature C-terminal domain (FhaB residues 1981 to 2471; Julio 2009), mouse monoclonal antibody that recognizes the AC domain of ACT (3D1; generated using ACT residues 373 to 399; Lee 1999; supply courtesy of F. Heath Damron), mouse monoclonal antibody that recognizes the RTX domain of ACT (9D4; generated using ACT residues 1156 to 1489; Hewlett 1989; supply courtesy of F. Heath Damron) or rabbit polyclonal antibody generated against the entire ACT protein. Corresponding α-rabbit and α-mouse IRDye secondary antibodies (LI-COR Biosciences) were used to detect proteins using a LI-COR Odyssey Classic Blot Imager (LI-COR Biosciences). *B. bronchiseptica* sample volumes were normalized based on optical density of the cultures.

To examine cell surface exoproteins by dot blot, bacteria were washed with PBS, and 100μL of 0.5OD_600_/mL bacterial suspension were spotted onto nitrocellulose membranes using a 96-well vacuum manifold. Membranes were probed using α-HA, α-FhaB, or polyclonal α-ACT antibodies and then by secondary antibodies listed above.

### Edman sequencing to identify SphB1-dependent cleavage site on ACT

The *B. bronchiseptica* RBX9 Δ*batB* strain was grown overnight in SS media supplemented with 25 μM Streptomycin. The supernatants were collected and 110 μL of a 1:1 mixture of culture supernatants and 2X Laemmli buffer were run on 5% SDS-PAGE at 60 mA for 2 hours. The proteins were transferred to PVDF membranes at 250 mA for 16 hours at 4°C. The membranes were stained with Coomassie Blue stain (5% acetic acid, 30% methanol, 1 g/L Coomassie R250). The band corresponding to ACT^C2^ was marked, the blot was destained in water to remove Coomassie, and the portion of the membrane containing ACT^C2^ was excised from the membrane and was submitted to the Stanford Protein and Nucleic Acid Facility (PAN) for Edman sequencing of the N terminal end of the ACT^C2^ polypeptide.

### Host cell intoxication assays

J774A.1 mouse-derived macrophage-like cell line was obtained from ATCC. CHO-K1 Chinese hamster ovary epithelial-like cell line and CHO-K1 producing human CR3 were supplied courtesy of Peter Sebo. Eukaryotic cells were grown in Dulbecco’s Modified Eagle Medium with high glucose and pyruvate (Thermo Fisher), supplemented with 10% fetal bovine serum (VWR) and 2 mM L-glutamine (Gibco). J774A.1 cells additionally were supplemented with 1% MEM Non-Essential Amino Acids (Gibco). To examine ACT intoxication of eukaryotic cells, medium was removed from 6-well plates of eukaryotes and replaced with bacteria-containing growth media. Plates were spun for 5 minutes at 500 g to bring bacteria into contact with the eukaryotic cells, then incubated at 37°C for 15 or 30 minutes. Cells were lysed with 0.1M HCl with 0.5% TritonX-100, and cell debris was pelleted by centrifugation at 21k xg for 10 minutes. Levels of cAMP in the supernatants were determined by competitive ELISA (ENZO).

## Acknowledgements

We would like to thank Dr. Peter Sebo for kindly providing the transgenic CHO cell line producing the human CR3 receptor and Dick Winant at the Stanford PAN Facility for Edman sequencing to identify the ACT^C2^ N terminus. We appreciate many productive conversations with the members of the Cotter lab regarding this work.

## Bibliography

1. Zaretzky FR, Gray MC, Hewlett EL. Mechanism of association of adenylate cyclase toxin with the surface of Bordetella pertussis: a role for toxin–filamentous haemagglutinin interaction. Mol Microbiol. 2002;45(6):1589–98.

2. Edwards JA, Groathouse NA, Boitano S. Bordetella bronchiseptica Adherence to Cilia Is Mediated by Multiple Adhesin Factors and Blocked by Surfactant Protein A. Infect Immun. 2005;73(6):3618–26.

3. Urisu A, Cowell JL, Manclark CR. Filamentous hemagglutinin has a major role in mediating adherence of Bordetella pertussis to human WiDr cells. Infect Immun. 1986;52(3):695–701.

4. Relman DA, Domenighini M, Tuomanen E, Rappuoli R, Falkow S. Filamentous hemagglutinin of Bordetella pertussis: nucleotide sequence and crucial role in adherence. Proc National Acad Sci. 1989;86(8):2637–41.

5. Cotter PA, Yuk MH, Mattoo S, Akerley BJ, Boschwitz J, Relman DA, et al. Filamentous Hemagglutinin of Bordetella bronchiseptica Is Required for Efficient Establishment of Tracheal Colonization. Infect Immun. 1998;66(12):5921–9.

6. Serra DO, Conover MS, Arnal L, Sloan GP, Rodriguez ME, Yantorno OM, et al. FHA-mediated cell-substrate and cell-cell adhesions are critical for Bordetella pertussis biofilm formation on abiotic surfaces and in the mouse nose and the trachea. Plos One. 2011;6(12):e28811.

7. Hoffman C, Eby J, Gray M, Damron FH, Melvin J, Cotter P, et al. Bordetella Adenylate Cyclase Toxin Interacts with Filamentous Hemagglutinin to Inhibit Biofilm Formation in vitro. Mol Microbiol. 2016;103(2):214–28.

8. Irie Y, Mattoo S, Yuk MH. The Bvg Virulence Control System Regulates Biofilm Formation in Bordetella bronchiseptica. J Bacteriol. 2004;186(17):5692–8.

9. Inatsuka CS, Julio SM, Cotter PA. Bordetella filamentous hemagglutinin plays a critical role in immunomodulation, suggesting a mechanism for host specificity. P Natl Acad Sci Usa. 2005;102(51):18578–83.

10. Henderson MW, Inatsuka CS, Sheets AJ, Williams CL, Benaron DJ, Donato GM, et al. Contribution of Bordetella filamentous hemagglutinin and adenylate cyclase toxin to suppression and evasion of interleukin-17-mediated inflammation. Infect Immun. 2012;80(6):2061–75.

11. Julio SM, Inatsuka CS, Mazar J, Dieterich C, Relman DA, Cotter PA. Natural-host animal models indicate functional interchangeability between the filamentous haemagglutinins of Bordetella pertussis and Bordetella bronchiseptica and reveal a role for the mature C-terminal domain, but not the RGD motif, during infection. Mol Microbiol. 2009;71(6):1574–90.

12. Melvin JA, Scheller EV, Noël CR, Cotter PA. New Insight into Filamentous Hemagglutinin Secretion Reveals a Role for Full-Length FhaB in Bordetella Virulence. Mbio. 2015;6(4):e01189–15.

13. Nash ZM, Cotter PA. Bordetella Filamentous Hemagglutinin, a Model for the Two-Partner Secretion Pathway. Microbiol Spectr. 2019;7(2).

14. Mazar J, Cotter PA. New insight into the molecular mechanisms of two-partner secretion. Trends Microbiol. 2007;15(11):508–15.

15. Jacob-Dubuisson F, Guérin J, Baelen S, Clantin B. Two-partner secretion: as simple as it sounds? Res Microbiol. 2013;164(6):583–95.

16. Chevalier N, Moser M, Koch HG, Schimz KL, Willery E, Locht C, et al. Membrane Targeting of a Bacterial Virulence Factor Harbouring an Extended Signal Peptide. J Mol Microb Biotech. 2005;8(1):7–18.

17. Lambert-Buisine C, Willery E, Locht C, Jacob-Dubuisson F. N-terminal characterization of the Bordetella pertussis filamentous haemagglutinin. Mol Microbiol. 1998;28(6):1283–93.

18. Clantin B, Hodak H, Willery E, Locht C, Jacob-Dubuisson F, Villeret V. The crystal structure of filamentous hemagglutinin secretion domain and its implications for the two-partner secretion pathway. P Natl Acad Sci Usa. 2004;101(16):6194–9.

19. Renauld-Mongénie G, Cornette J, Mielcarek N, Menozzi FD, Locht C. Distinct roles of the N-terminal and C-terminal precursor domains in the biogenesis of the Bordetella pertussis filamentous hemagglutinin. J Bacteriol. 1996;178(4):1053–60.

20. Jacob-Dubuisson F, Buisine C, Willery E, Renauld-Mongénie G, Locht C. Lack of functional complementation between Bordetella pertussis filamentous hemagglutinin and Proteus mirabilis HpmA hemolysin secretion machineries. J Bacteriol. 1997;179(3):775–83.

21. Mazar J, Cotter PA. Topology and maturation of filamentous haemagglutinin suggest a new model for two-partner secretion. Mol Microbiol. 2006;62(3):641–54.

22. Noël CR, Mazar J, Melvin JA, Sexton JA, Cotter PA. The prodomain of the Bordetella two-partner secretion pathway protein FhaB remains intracellular yet affects the conformation of the mature C-terminal domain. Mol Microbiol. 2012;86(4):988–1006.

23. Johnson RM, Nash ZM, Dedloff MR, Shook JC, Cotter PA. DegP Initiates Regulated Processing of Filamentous Hemagglutinin in Bordetella bronchiseptica. Mbio. 2021;12(3):e01465–21.

24. Nash ZM, Cotter PA. Regulated, sequential processing by multiple proteases is required for proper maturation and release of Bordetella filamentous hemagglutinin. Mol Microbiol. 2019;112(3):820–36.

25. Coutte L, Antoine R, Drobecq H, Locht C, Jacob-Dubuisson F. Subtilisin-like autotransporter serves as maturation protease in a bacterial secretion pathway. Embo J. 2001;20(18):5040–8.

26. Wolff J, Cook GH, Goldhammer AR, Berkowitz SA. Calmodulin activates prokaryotic adenylate cyclase. Proc Natl Acad Sci. 1980;77(7):3841–4.

27. Hewlett EL, Urban MA, Manclark CR, Wolff J. Extracytoplasmic adenylate cyclase of Bordetella pertussis. Proc National Acad Sci. 1976;73(6):1926–30.

28. Hackett M, Guo L, Shabanowitz J, Hunt DF, Hewlett EL. Internal Lysine Palmitoylation in Adenylate Cyclase Toxin from Bordetella pertussis. Science. 1994;266(5184):433–5.

29. Barry EM, Weiss AA, Ehrmann IE, Gray MC, Hewlett EL, Goodwin MS. Bordetella pertussis adenylate cyclase toxin and hemolytic activities require a second gene, cyaC, for activation. J Bacteriol. 1991;173(2):720–6.

30. Glaser P, Sakamoto H, Bellalou J, Ullmann A, Danchin A. Secretion of cyclolysin, the calmodulin-sensitive adenylate cyclase-haemolysin bifunctional protein of Bordetella pertussis. Embo J. 1988;7(12):3997–4004.

31. Šebo P, Ladant D. Repeat sequences in the Bordetella pertussis adenylate cyclase toxin can be recognized as alternative carboxy-proximal secretion signals by the Escherichia coliα-haemolysin translocator. Mol Microbiol. 1993;9(5):999–1009.

32. Osicka R, Osickova A, Hasan S, Bumba L, Cerny J, Sebo P. Bordetella adenylate cyclase toxin is a unique ligand of the integrin complement receptor 3. Elife. 2015;4:e10766.

33. Cotter PA, Miller JF. A mutation in the Bordetella bronchiseptica bvgS gene results in reduced virulence and increased resistance to starvation, and identifies a new class of Bvg-regulated antigens. Mol Microbiol. 1997;24(4):671–85.

34. Bumba L, Masin J, Macek P, Wald T, Motlova L, Bibova I, et al. Calcium-Driven Folding of RTX Domain β-Rolls Ratchets Translocation of RTX Proteins through Type I Secretion Ducts. Mol Cell. 2016;62(1):47–62.

35. Stainer DW, Scholte MJ. A Simple Chemically Defined Medium for the Production of Phase I Bordetella pertussis. Microbiology+. 1970;63(2):211–20.

36. Lee SJ, Gray MC, Guo L, Sebo P, Hewlett EL. Epitope Mapping of Monoclonal Antibodies against Bordetella pertussis Adenylate Cyclase Toxin. Infect Immun. 1999;67(5):2090–5.

37. Guermonprez P, Khelef N, Blouin E, Rieu P, Ricciardi-Castagnoli P, Guiso N, et al. The adenylate cyclase toxin of Bordetella pertussis binds to target cells via the alpha(M)beta(2) integrin (CD11b/CD18). J Exp Medicine. 2001;193(9):1035–44.

38. Gray MC, Donato GM, Jones FR, Kim T, Hewlett EL. Newly secreted adenylate cyclase toxin is responsible for intoxication of target cells by Bordetella pertussis: Active secretion of adenylate cyclase toxin. Mol Microbiol. 2004;53(6):1709–19.

39. Confer D, Eaton J. Phagocyte impotence caused by an invasive bacterial adenylate cyclase. Science. 1982;217(4563):948–50.

40. Utsumi S, Sonoda S, Imagawa T, Kanoh M. Polymorphonuclear leukocyte-inhibitory factor of Bordetella pertussis. I. Extraction and partial purification of phagocytosis- and chemotaxis-inhibitory activities. Biken J. 1978;21(4):121– 35.

41. Hewlett EL, Gordon VM, McCaffery JD, Sutherland WM, Gray MC. Adenylate cyclase toxin from Bordetella pertussis Identification and purification of the holotoxin molecule*. J Biol Chem. 1989;264(32):19379–84.

42. Šebo P, Glaser P, Sakamoto H, Ullmann A. High-level synthesis of active adenylate cyclase toxin of Bordetella pertussis in a reconstructed Escherichia coli system. Gene. 1991;104(1):19–24.

43. Ladant D, Michelson S, Sarfati R, Gilles AM, Predeleanu R, Bârzu O. Characterization of the Calmodulin-binding and of the Catalytic Domains of Bordetella pertussis Adenylate Cyclase. J Biol Chem. 1989;264(7):4015–20.

44. MUNIER H, BOUHSS A, GILLES A, KRIN E, GLASER P, DANCHIN A, et al. Structural flexibility of the calmodulin-binding locus in Bordetella pertussis adenylate cyclase. Eur J Biochem. 1993;217(2):581–6.

45. Karimova G, Pidoux J, Ullmann A, Ladant D. A bacterial two-hybrid system based on a reconstituted signal transduction pathway. Proc Natl Acad Sci. 1998;95(10):5752–6.

46. Ruhe ZC, Subramanian P, Song K, Nguyen JY, Stevens TA, Low DA, et al. Programmed Secretion Arrest and Receptor-Triggered Toxin Export during Antibacterial Contact-Dependent Growth Inhibition. Cell. 2018;175(4):921–933.e14.

47. Zekarias B, Mattoo S, Worby C, Lehmann J, Rosenbusch RF, Corbeil LB. Histophilus somni IbpA DR2/Fic in Virulence and Immunoprotection at the Natural Host Alveolar Epithelial Barrier. Infect Immun. 2010;78(5):1850–8.

48. Allen JP, Ozer EA, Minasov G, Shuvalova L, Kiryukhina O, Satchell KJF, et al. A comparative genomics approach identifies contact-dependent growth inhibition as a virulence determinant. P Natl Acad Sci Usa. 2020;117(12):6811–21.

